## Supplementary Table 3 for "Taste shaped the use of botanical drugs"

**Supplementary file 3. Codes and the short descriptions of the 22 chemosensory qualities to represent tastes and flavours. Codes are those used in Supplementary files 2,5 and 6; short descriptions match those used in Figure 1.**

1.	**AROM**:	Aromatic

2.	**ASTR**:	Astringent

3.	**BALS**:	Balsamic

4.	**BITT**:	Bitter

5.	**BURN**:	Burning/Hot

6.	**FRES:**	Fresh/Cooling

7.	**FRUI:**	Fruity

8.	**HERB**:	Herby/Leafy

9.	**MUCI:**	Mucilaginous

10.	**MUSK**:	Musky

11.	**NUTT**:	Nutty

12.	**SALT**:	Salty

13.	**SMOK**:	Smoky

14.	**SOAP**:	Soapy

15.	**SOUR**:	Sour/Acidic

16.	**STAR**:	Starchy

17.	**STGI**:	Stinging/Mustardy

18.	**STNK**:	Stinky

19.	**STRW**:	Straw-like

20.	**SWEE**:	Sweet

21.	**WARM**:	Warming

22.	**WOOD**:	Woody
