## Supplementary Table 4 for "Taste shaped the use of botanical drugs"

**Supplementary file 4. Codes and the short descriptions of the therapeutic uses as used in Supplementary files 1,5 and 6; short descriptions match those used in Figure 1. Notes present a fuller description of the therapeutic uses as in *DMM* (*ex* Matthioli, 1568, Supplementary file 1). The number of use records for each therapeutic use is given.**

| **Category of use code and short description** | **Notes** | **Use records** |
| --- | --- | --- |
| **ANDR:** Andrology | Improve fertility, sterilization, contraceptives | 12 |
| **ANTI:** Antidotes | Bites and stings from venomous animals incl. snakes, arthropods, sea animals and remedies for envenomations, intoxications and food poisonings, applied internally | 230 |
| **CARD:** Cardiac problems | Cardiac and precordial chest pain | 7 |
| **EARS:** Ears | Tinnitus, deafness | 22 |
| **EYES:** Eyes | Lacrimal fistula, red, watery and tired eyes, blurred vision, hordeolum, cataract | 178 |
| **FEVE:** Fever | Periodic, tertian and quartan fever, shivering (including malaria) | 40 |
| **FOOD:** Food | Good for the stomach, nutritious, bread making, condiment, diet (also used for diet) | 109 |
| **GA.DI:** Gastric - Diarrheal conditions | Diarrhoea and dysentery | 190 |
| **GA.FU:** Gastric - Function | Appetizers, digestives, abdominal and intestinal pain, promote digestion, heartburn, heat in the stomach, weak stomach, stomach-ache, stomach problems, hiccup, colic, flatulence, nausea, vomiting, hematemesis, intestinal ruptures, anal prolapse | 230 |
| **GA.LI:** Gastric - Liver, jaundice | Liver problems and jaundice | 87 |
| **GA.PL:** Gastric – Purgatives | Soften and purge the belly, purge the stomach, laxatives, emetics, drive out intestinal worms, kill intestinal worms | 125 |
| **GOUT:** Gout | Gout | 31 |
| **GY.AE:** Gynaecology - Abortion, menses | Abortifacients, uterotonics emmenagogues, inducing delivery | 272 |
| **GY.BR:** Gynaecology – Breast | Promote lactation, induration and inflammation of the breast (mastitis) | 39 |
| **GY.OT:** Gynaecology – Other | Contraceptives, promote conception, inflamed vagina, vaginal ulcers, prevent the growth of the breasts, stop lactation | 39 |
| **GY.UT:** Gynaecology - Uterine | Uterus | 91 |
| **GY.VD:** Gynaecology - Vaginal discharge | Staunch vaginal discharge (leucorrhoea, menstrual discharge) | 65 |
| **HMM:** Humoral | Humoral management: purge yellow and black bile, thick humours, phlegm (watery humours), for the choleric | 46 |
| **LIBI:** Libido | Aphrodisiacs and anaphrodisiacs, prevent sexual dreams | 33 |
| **MOUTH:** Oral cavity | Problems of the teeth, gums and oral cavity: ulcers, sores, gingivitis, loose teeth hoarseness; hygiene: mouthwash, cleaning teeth | 85 |
| **MU.EX:** Musculoskeletal – External | Bone fractures, dislocations, injured muscles, joints and tendons, spasms, lower back pain, all sorts of bruises, for the ruptured and the spastics | 81 |
| **MU.IN:** Musculoskeletal -Internal | Bone fractures, dislocations, injured muscles, joints and tendons, spasms, lower back pain, all sorts of bruises, for the ruptured and the spastics | 58 |
| **NS.PA:** Neurological – Pain | Headache, toothache, earache, stitch, unspecific pain | 185 |
| **NS.PS:** Neurological - Psychiatric | Soporifics, against melancholy and frenzy, inducing sleep and against fatigue | 68 |
| **NS.SC:** Neurological – Sciatica | Sciatica | 51 |
| **NS.VA:** Neurological - Various incl. epilepsy | Epilepsy, tremor, paralysis, hangover, numbness, vertigo | 53 |
| **NOSE:** Nose | Epistaxis, polyps, purge the head from phlegm, provoke sneezing | 28 |
| **PARA:** Parasites | Lice, fleas, scabies, worms in the ear | 57 |
| **POIS:** Poisons | To kill vertebrates and invertebrates incl. hunting purposes and fish poisons | 26 |
| **REPE:** Repellents | Repelling snakes, scorpions, wasps, mosquitoes, moths, fleas | 36 |
| **RE.BD:** Respiratory - Breathing difficulties | Dyspnoea, asthma | 62 |
| **RE.CO:** Respiratory – Cough | Catarrhs, chronic, dry and bloody cough, inflamed throat, purge chest, lungs and throat | 184 |
| **RE.OT:** Respiratory – Other | Tuberculosis, pneumonia, pleurisy, unspecific chest complaints, lost voice, lung defects, tonsillitis, diphtheria | 56 |
| **SK.AB:** Skin - Animal bites | Treatments for bites and stings of venomous and non-venomous animals, applied externally | 92 |
| **SK.CO:** Skin - Cosmetics | Hair loss, hair colouring, dandruff, nails, vitiligo, eschars, scars, freckles, clean and tighten skin, against perspiration, armpit odour, perfumes, cosmetics | 193 |
| **SK.IF:** Skin - Infections | Inflamed wounds, erysipelas, impetigo, lepra, abscesses, furuncles, carbuncles, pustules, gangrene, paronychia, fistula, scrofula | 265 |
| **SK.IN:** Skin - Inflammation | Pruritus, dermatitis, blisters, boils, eruptions, burns, sunburns, chilblains | 158 |
| **SK.OT:** Skin - Other | Indurations and excrescences, tumours, warts, callus, swellings | 106 |
| **SK.UL:** Skin - Topical ulcers | Filthy, creeping and chronic ulcers | 180 |
| **SK.WO:** Skin - Wounds | Bleeding wounds, fissures, abrasions, splinters | 86 |
| **SPLE:** Spleen | Swollen spleen, indurations | 63 |
| **UR.CA**: Urinary - Calculus | Renal and vesical calculi | 40 |
| **UR.DI:** Urinary - Diuretics | Dropsy, oedema | 224 |
| **UR.OT:** Urinary - Other | Kidney and bladder problems, urinary retention, difficult urination | 105 |
| **VAR:** Various | Enchantment, reduce weight, bleedings, cold, unspecified abscesses, weak limbs | 33 |
| **VASC:** Vascular | Haemorrhoids and varices | 12 |
